# Tail length in male versus female fox squirrels (*Sciurus niger*)

**DOI:** 10.1101/2023.08.08.552536

**Authors:** Hannah K. Nichols, Shaylee K. Smith, Valerie M. Eddington, Adrienne Calistri-Yeh, Laura N. Kloepper, Vanessa K Hilliard Young

**Affiliations:** Department of Biology, Saint Mary’s College, Notre Dame, IN 46556 USA; School for Environment and Sustainability, University of Michigan, Ann Arbor, MI 48109 USA; Department of Biological Sciences and Center for Acoustics Research and Education, University of New Hampshire, NH 03824 USA

**Keywords:** arboreal, locomotion, morphology, reproduction, squirrel

## Abstract

**Background:** Arboreal mammals rely on their tails to aid in balance while maneuvering complex habitats. Females experience additional challenges to locomotion due to reproductive demands including altered body mass and/or body shape, which leads to shifts in center of mass. Without compensation, this may increase the risk of losing balance and falling out of trees. We tested the hypothesis that female squirrels have longer tails than males to offset shifts in center of mass that may result from pregnancy.

**Results:** Morphological data were collected from 57 fox squirrels (*Sciurus niger*) in northern Indiana in summer 2019 and 2021. Although our initial t-test analysis of relative tail length (RTL) showed that female squirrels had longer tails than males (*p* = 0.02), a subsequent ANCOVA that controlled for effect of body length indicated no significant effect of sex on tail length (*p* = 0.42).

**Conclusions:** The results of this study demonstrate the potential impacts of different analysis methods on overall understanding of organismal functional morphology and are an important addition to the literature on tail form and function, which remains poorly understood compared to other appendages.

## BACKGROUND

Arboreal habitats consist of complex environments, often including substrates with varying thicknesses and stability (1, 2). Maneuvering in these habitats presents specific challenges to arboreal animals, especially with regard to balance. Such challenges include but are not limited to, proper foot placement, grasping ability, and tail usage (1). Miscalculations or difficulty navigating these challenges can cause an animal to lose its balance, which can lead to injury or death and ultimately decrease individual fitness. To successfully move about their habitats, arboreal animals present behavioral adjustments and morphological adaptations in response to instability during locomotion (2, 3, 4, 5, 6, 7). When subjected to strictly arboreal environments, laboratory mice displayed behaviors of tail usage and pedal grasping to adapt to the new environment (2). Quadrupedal monkeys were also observed to exhibit dramatic tail whip motions during periods of instability during arboreal locomotion (3). In addition to behavioral adaptations, many arboreal animals have morphological adaptations to promote stability in their environment (4, 5, 6, 7). For example, European red squirrels (*Sciurus vulgaris L*.) have longer forelimbs, shoulder muscles with increased flexing ability, and shorter tibia and hindlimbs to aid in clawed locomotion while on narrow terminal branches (6). Likewise, for arboreal mammals, navigating arboreal substrates selected for longer tails (7). A longer tail functions better as a counterweight than a short tail, which can facilitate balance (7). Behavioral adjustments along with morphological adaptations allow arboreal animals to locomote successfully in their environment.

Reproductive and maternal demands can have substantial negative impacts on females, including reducing foraging efficiency, predator avoidance, and other essential activities by decreasing locomotor abilities (8, 9, 10, 11, 12, 13). In general, larger female mammals tend to have better reproductive outputs (11), but with the increased size comes significant costs. In particular, body size and weight while gravid can greatly impact locomotor abilities (12, 13). For example, pregnant bottlenose dolphins (*Tursiops truncatus*) show a decrease in locomotor performance and an increase in drag resulting from changes in morphology due to the increased load associated with pregnancy (13). Similarly, wild brown anoles (*Anolis sagrei*) show decreased locomotor abilities while gravid, also due to increased load in addition to restraints on energy allocation (12). Likewise, when compared to non-pregnant female counterparts, pregnant garter snakes (*Thamnophis marcianus*) had significantly lower locomotor abilities (14). Increased body mass and negative effects on energy allocation that are associated with pregnancy decrease locomotor performance and abilities across multiple species.

Considering the extensive locomotor demands arboreal animals are subjected to in regard to balance, pregnancy can have detrimental effects on stability while navigating their habitat (4, 15, 16). When gravid, an increase in body mass shifts the center of mass, resulting in a reduced ability to balance (15). A shifting center of mass coupled with an unsteady and complex habitat to maneuver in adds locomotor challenges for arboreal females. This can lead to selection pressures favoring certain conditions for females that would benefit their locomotor abilities while gravid (4; 12, 16). Selection pressures on females due to locomotive challenges associated with gravidity have been identified in other species. For instance, anoles are believed to have evolved a single-egg clutch to help increase locomotor abilities while gravid (12). The selection pressures faced by arboreal squirrels could also favor certain conditions in females, such as long tails to help maintain center of mass.

Tails have been found to be both static and dynamic tools that aid in balance for arboreal quadrupeds (3, 6, 17). Arboreal animals with shorter digits and claws rely heavily on their tails, using them as dynamic stabilizers versus counterweights to improve stability (6). Consequently, lizards (*Anolis carolinensis*) that experience caudal autotomy have a reduction in jumping performance and can unintentionally rotate midair without their tails (18). This suggests that tails are necessary structures for controlling whole-body rotation and may be valuable for effective navigation in arboreal habitats.

Significant correlations between tail morphology and locomotion have been observed (3, 4, 6, 7, 17). In general, for arboreal animals, longer tails are better suited for arboreal locomotion (3, 4, 6, 7). For example, ground squirrels that do not engage in arboreal locomotion have shorter tails compared to tree squirrels (4). Longer tails can be indicative of not only a squirrel’s habitat but also the complexity of navigating their habitat. Another important factor to note is the sexual dimorphism observed in squirrels (4, 19, 20). A study by Pasch and Koprowski (20) reported that female Chiricahua fox squirrels (*Sciurus nayaritensis chiricahuae*) experienced seasonal body mass fluctuations, unlike their male counterparts. It was also noted that males and females only differed in body mass during the winter and spring, with females being heavier than males, coinciding with parturition (20). Fluctuations and an increased body mass could require a longer tail to offset the locomotor challenges these characteristics create. Indeed, Hayssen (4) found that female squirrels from several genera (subfamilies: Protoxerini, Pteromyini, Callosciurinae, Ratufinae, Sciurillinae, and Sciurini) tended to have longer tails than males. Although Hayssen (4) includes the genera in the group Sciurini, it is unclear what were the sample size and the tail length data for fox squirrels (*Sciurus niger*), the largest arboreal squirrel species native to North America. Therefore, we sought to assess tail length in male and female fox squirrels, to determine whether tail length patterns observed in other taxa extended to this species. Given results from prior studies (4, 19, 20) we hypothesized that female fox squirrel tail lengths will be longer than male tail lengths relative to body length.

## RESULTS

Data were collected from 57 individual adult fox squirrels: 33 males and 24 females. In total, 18 of the 57 squirrels were recaptured and their morphological measurements were averaged.

Tail length and body length showed normal distribution based on results from a Shapiro-Wilks normality test (W = 0.98, *p* = 0.62 and W = 0.98, *p* = 0.39, respectively). The average female relative tail length (RTL; 0.80 ± 0.10) was 8.1% longer than the average male RTL (0.74 ± 0.11) (Fig. 1). There was a significant difference in RTL between males and females (*t*(55) = 2.05, *p* = 0.02).

**Figure 1.**
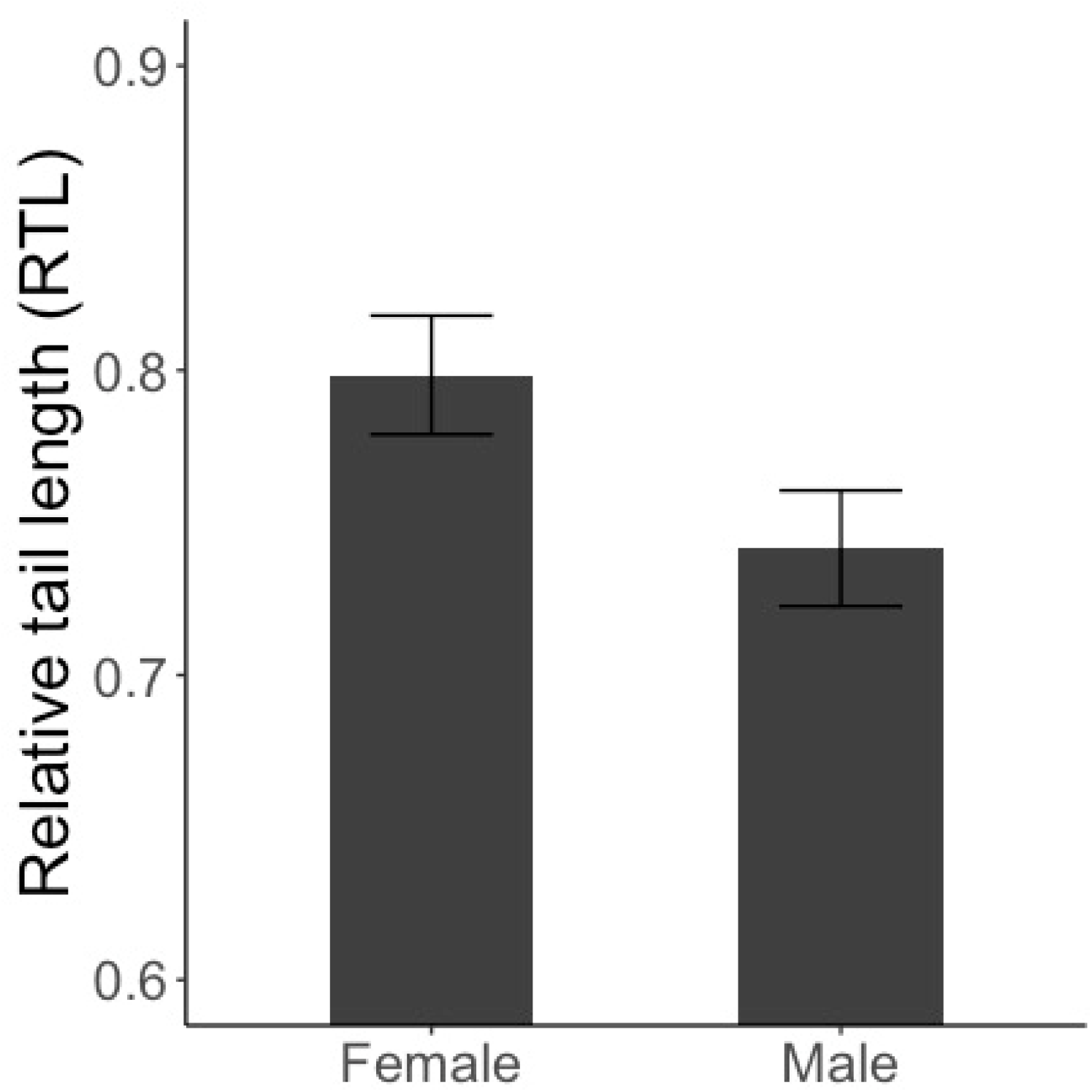
Average relative tail length (RTL) of female versus male fox squirrels (n = 24 and n = 33, respectively). Average RTL for females was 0.80 ± 0.10; for males average RTL was 0.74 ± 0.11. Error bars indicate standard error.

The covariate, body length, and sex are independent (F_(1,55)_ = 2.41, *p* = 0.13). The variances among the groups are equal (F_1,55)_ = 1.73, *p* = 0.19). The covariate, body length, was not significantly related to tail length (F_(1,57)_ = 1.09, *p* = 0.30). There was no significant effect of sex on tail length after controlling for body length (F_(1,57)_ = 0.44, *p* = 0.51) (Fig. 2).

**Figure 2.**
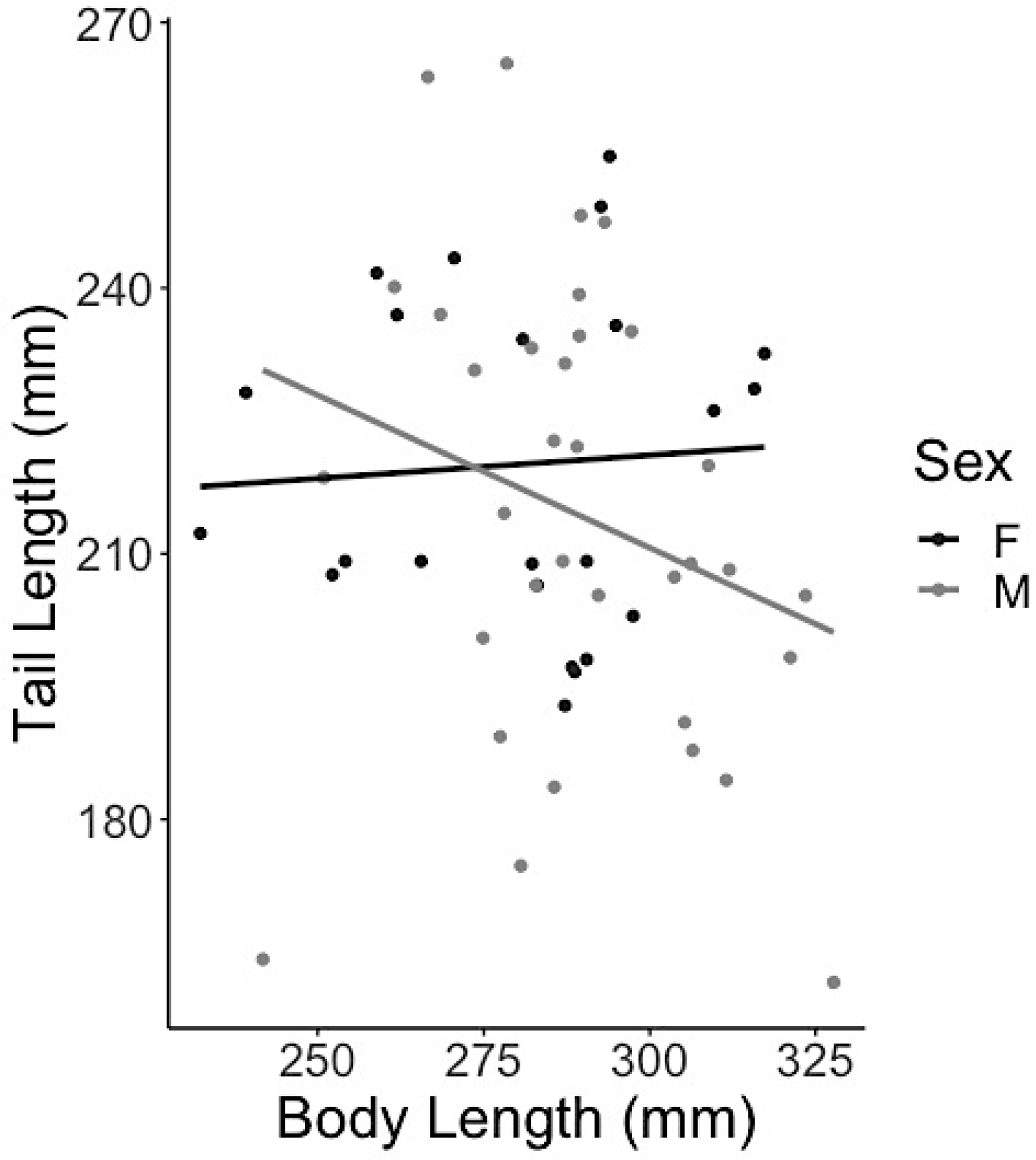
Tail length vs body length in male (grey) and female (black) fox squirrels with lines of best fit.

## DISCUSSION

The purpose of this study was to compare the relative tail lengths of female fox squirrels to their male counterparts. Our results do not definitively support the hypothesis that female fox squirrels have significantly longer tails compared to males.

When following the methods of Hayssen (4) our results were consistent with their analysis of squirrel tail lengths. Data collected from museum specimens demonstrated that female tree squirrels had tails that were 1.8% longer than those of males and 3.8% longer for genera in the group Sciurini (4). Our results indicate a larger difference between male and female tail lengths for fox squirrels (8.1% greater RTL; Fig. 1) than those of Hayssen (4) while using similar methods.

When controlling for body length, however, our results a non-significant difference in tail length. Hayssen (4) calculated relative tail length by dividing tail length by body length and conducted a t-test; our study used this method but also included a post hoc analysis of covariance (ANCOVA) to assess tail length relative to body length. When controlling for body length, we found that sex did not affect tail length (Fig. 2). These results could be explained by the research of Mincer and Russo (7), which suggests arboreal tail lengths scale to body length with negative allometry. Females tend to be smaller than their male counterparts, so considering this difference in overall body size and the negative allometric relationship between tail length and body length could account for our results.

The ANCOVA allowed us to check for a consistent difference in tail length between sexes across all body lengths. Aside from this singular study by Hayssen (4), there are no other published sources that have looked specifically at tail lengths in arboreal squirrels. Considering the difference in the results of our study using these two analysis methods, further research is necessary for evaluating and understanding the functional morphology of tails in arboreal taxa. Possible directions for future research could examine tail use by pregnant squirrels, tail use in male and female squirrels during the breeding season, and in female squirrels while not pregnant. Such data could provide valuable insights to behavioral differences in tail use relative to sex and/or reproductive state in arboreal squirrels. The importance of future research is exacerbated by the literature that points to longer tails being advantageous to arboreal females.

Arboreal animals, such as fox squirrels, rely on their tails to act as a balancing mechanism during locomotion (3, 6, 17). Females, due to reproductive demands, face additional challenges to their balance (8, 9, 10; 11, 12). The additional challenges due to gestation and related body mass fluctuations suggest for the selection of a longer tail compared to males (16). For example, ‘flying lizards’ (*Draco melanopogon*) have been found to have larger body sizes and longer tails to compensate for the effects of pregnancy on gliding locomotion (9). Similarly, southern flying squirrels (*Glaucomys volans*) were found to have larger body sizes to accommodate for the added mass during pregnancy which affects wing-loading (21). Evidence of evolutionarily favored morphological adaptations for different taxa suggests that longer tails in arboreal squirrels could be influenced by reproductive demands (16).

Although locomotor mechanics may be an important driver of tail morphology in arboreal taxa, there are likely additional factors related to tail function that may affect tail length. For example, tails are used for pre- and post-copulatory communication (22) and communicating in aggressive encounters with conspecifics (23). Furthermore, in the presence of snakes, squirrels tail flag as a method of strike avoidance behavior (24). A longer tail may be beneficial for tail flagging as the movement would be more pronounced and give individuals more time to avoid strikes. This additional time could be particularly advantageous for females during pregnancy, as their locomotor abilities are decreased. There may also be evolutionary advantages or sexual selection for shorter tails in males; however, we are aware of no research in support of these hypotheses.

## CONCLUSIONS

Tails are a common morphological feature of vertebrate animals, however the evolutionary importance of tails is poorly understood compared to other appendages such as forelimbs (25). The current research on tail form and function across all species is insufficient to be able to make complex analyses and observations from a comparative and broader perspective (25). Despite the lack of research, one important correlation that has been found is between tail length and arboreality (26, 16). Longer tails are selected for arboreal habitats due to their stabilizing abilities (3, 4, 6, 7, 16, 27). Tails, specifically longer ones, can also become a complication during predator escape. Longer tails could increase drag while running or allow for a predator to grab onto at further distances (4). Given these implications, there are many tradeoffs for longer tails in arboreal taxa. Our results can provide insight into the tradeoffs and evolutionary implications for arboreal squirrels and tail lengths; however, future work on arboreal maneuverability at different stages of the reproductive cycle may provide additional insights to tail usage in squirrels and other taxa.

## METHODS

Data were collected from Eastern fox squirrels (*Sciurus niger*) on the Saint Mary’s College campus (Notre Dame, IN) for nine weeks during May-July 2021. Eight live animal traps (collapsible squirrel traps #203, Tomahawk Live Trap, Hazelhurst, WI, USA) were baited with an apple slice, peanut butter, and sunflower seeds, and covered with a brown canvas to provide shade and reduce stress. The traps were set at 0900 Monday through Thursday and examined every three hours at 1200 and 1500. Traps were removed from the field daily at 1500. When captured, squirrels were ear tagged (self-piercing ear tags #1005-3, National Band and Tag Company, Newport, KY, USA) for identification purposes, and the following morphological measurements were recorded: body mass, body length, and tail length. Tails were palpated during measurement to detect truncated tails; visibly shortened tails were also noted. Tail length was measured from the base of the tail to the last caudal vertebra and body length was measured from the nose to the base of the tail. Sex and reproductive state were also recorded, if discernible. Squirrels were handled using canvas cones proposed by Koprowski (28). This handling device was chosen as it is effective in minimizing handling shock on non-anesthetized squirrels, eliminates the costs associated with using anesthesia, and reduces bite risk for the researchers (28). After data collection, squirrel ears were carefully exposed and self-piercing ear tags were applied to the middle of the ear, using a self-piercing tag applicator (self-piercing ear tags #1005-3, National Band and Tag Company, Newport, KY, USA). Squirrels were then released at the site of capture. All work was conducted in accordance with IACUC-approved methods (Saint Mary’s IACUC Protocol Number: 2020-007).

Data collected in summer 2019 provided an additional 31 samples, which were pooled with the data collected from the 2021 trapping season (N_2021_ = 26; N_total_ = 57). Recaptured squirrels were identified by ear tags and remeasured. Morphological measurements for squirrels caught more than once were averaged to obtain a more accurate measurement, accounting for measuring error; individuals with truncated tails were excluded from data analysis. Morphological measurements for recaptures was highly consistent across samples, with measurements varying by less than 2cm. Body length and tail length measurements were used for data analysis. A Shapiro-Wilks normality test was run on both variables of tail length and body length to confirm the measurements were normally distributed. Two different types of analyses were performed based on the methods of Hayssen (4) and Mincer and Russo (7).

Relative tail length to body length (RTL) was used to account for size variation between sexes. To calculate RTL, tail length (mm) was divided by body length (mm) (4). A two-sample, one-tailed *t*-test was conducted to compare the RTL of male and female squirrels. Finally, to compare the average RTL between males and females, male RTL was subtracted from female RTL and then divided by male RTL. To analyze the impact of sex on tail length while controlling for body length, a post hoc analysis of covariance was conducted (ANCOVA; 7). An ANOVA and Levene’s Test were run to verify independence and homogeneity of variance prior to the analysis of covariance. All calculations were conducted in R studio v.4.1.0 (29).

## LIST OF ABBREVIATIONS

RTL: Relative Tail Length
IACUC: Institutional Animal Care and Use Committee
ANCOVA: Analysis of Covariance

## DECLARATIONS

### Ethics approval and consent to participate

The ethical approval was provided by the Saint Mary’s College Institutional Animal Care and Use Committee (IACUC). All work was conducted in accordance with IACUC-approved methods (Saint Mary’s IACUC Protocol Number: 2020-007).

### Consent for publication

Not applicable.

### Availability of data and materials

The datasets used and/or analyzed during the current study are available from the corresponding author on reasonable request.

### Authors’ contributions

Study design: HKN, SKS, and VKHY; Data collection: HKN, VME, ACY, LNK, and VKHY; Data Analysis: HKN, SKS, and VKHY; Manuscript preparation: HKN, SKS, VME, ACY, LNK, and VKHY

## Acknowledgments

The authors wish to thank The Sisters of the Holy Cross for access to their land and Saint Mary’s College alumnae Hannah Gams and Kamryn Yerga for data collection. This project was funded by the Saint Mary’s College Marjorie Neuhoff Summer Science Communities Grant for Undergraduate Research and comprised the senior thesis of Hannah Nichols.

## REFERENCES

1. Preuschoft H. What does “arboreal locomotion” mean exactly and what are the relationships between “climbing”, environment and morphology? Zeitschrift für Morphologie und Anthropologie. 2002;83:171–88.

2. Byron C, Kunz H, Matuszek H, Lewis S, Van Valkinburgh D. Rudimentary pedal grasping in mice and implications for terminal branch arboreal quadrupedalism. Journal of Morphology. 2011;272:230–40.

3. Larson SG, Stern JT, Jr. Maintenance of above-branch balance during primate arboreal quadrupedalism: Coordinated use of forearm rotators and tail motion. American Journal of Physical Anthropology. 2006;129:71–81.

4. Hayssen V. Patterns of body and tail length and body mass in Sciuridae. Journal of Mammalogy. 2008;89:852–73.

5. Youlatos D, Samaras A. Arboreal locomotor and postural behavior in European red squirrels (Sciurus vulgaris L.) in northern Greece. Journal of Ethology. 2011;29:235–42.

6. Young JW, Russo GA, Fellmann CD, Thatikunta MA, Chadwell BA. Tail function during arboreal quadrupedalism in squirrel monkeys (Saimiri boliviensis) and tamarins (Saguinus oedipus). Journal of Experimental Zoology. 2015;323A:556-66.

7. Mincer ST, Russo GA. Substrate use drives the macroevolution of mammalian tail length diversity. Proceedings of the Royal Society B. 2020;287:20192885.

8. Altmann J, Samuels A. Costs of maternal care: infant-carrying in baboons. Behavioral Ecology and Sociobiology. 1992;29:391–8.

9. Shine R, Keogh S, Doughty P, Giragossyan H. 1998. Costs of reproduction and the evolution of sexual dimorphism in a ‘flying lizard’ Draco melanopogon (Agamidae). Journal of Zoology. 1998;246:203–13.

10. McLean JA, Speakman JR. Effects of body mass and reproduction on the basal metabolic rate of brown long-eared bats (Plecotus auritus). Physiological and Biochemical Zoology. 2000;73:112–21

11. Fokidis HB, Risch TS. The burden of motherhood: gliding locomotion in mammals influences maternal reproductive investment. Journal of Mammalogy. 2008a;89:617–25.

12. Cox RM, Calsbeek R. Severe costs of reproduction persist in Anolis lizards despite the evolution of a single-egg clutch. Evolution. 2009;64:1321–30.

13. Noren SR, Redfern JV, Edwards EF. Pregnancy is a drag: hydrodynamics, kinematics and performance in pre-and post-parturition bottlenose dolphins (Tursiops truncatus). The Journal of Experimental Biology. 2011;214:4151–9.

14. Seigel RA, Huggins MM, Ford NB. Reduction in locomotor ability as a cost of reproduction in gravid snakes. Oecologia (Berlin). 1987;73:481–5.

15. Foti T, Davids JR, Bagley, A. A biomechanical analysis of gait during pregnancy. Journal of Bone and Joint Surgery. 2000;82:625–32.

16. Smith SK, Young VKH. Balancing on a limb: effects of gravidity on locomotion in arboreal, limited vertebrates. Integrative and Comparative Biology. 2021;61:573–8.

17. Dunbar DC, Badam GL. Locomotion and posture during terminal branch feeding. International Journal of Primatology. 2000;421:649–69.

18. Gillis GB, Bonvini LA, Irschick DJ. Losing stability: tail loss and jumping in the arboreal lizard Anolis carolinensis. Journal of Experimental Biology. 2009;212:604–9.

19. Holekamp KE, Nunes S. Seasonal variation in body weight, fat, and behavior of California ground squirrels (Spermophilus beecheyi). Canadian Journal of Zoology. 1989;67:1425–33.

20. Pasch BS, Koprowski JL. Annual cycles in body mass and reproduction of Chiricahua fox squirrels (Sciurus nayaritensis chiricahuae). The Southwestern Naturalist. 2006;51:531–5.

21. Fokidis HB, Risch TS. Does gliding when pregnant select for larger females? Journal of Zoology. 2008b;275:237–44.

22. McCloskey RJ, Shaw KC. Copulatory Behavior of the Fox Squirrel. Journal of Mammology. 1977;58:663–665.

23. Pardo, MA. Eastern gray squirrels (Sciurus carolinensis) communicate with the positions of their tails in an agonistic context. The American Midland Naturalist. 2014; 172:359–365.

24. Putman BJ, Clark RW. The fear of unseen predators: ground squirrel tail flagging in the absence of snakes signals vigilance. Behavioral Ecology. 2015;26:185–93.

25. Schwaner MJ, Hsieh ST, Swalla BJ, McGowan CP. An introduction to an evolutionary tail: EvoDevo, structure, and function of post-anal appendages. Integrative and Comparative Biology. 2021;61:352–7.

26. Sheehy CM, Albert JS, Lillywhite HB. The evolution of tail length in snakes associated with different gravitational environments, Functional Ecology. 2016;30:244–54.

27. Young JW, Chadwell BA, Dunham NT, McNamara A, Phelps T, Hieronymus T, Shapiro LJ. 2021. The stabilizing function of the tail during arboreal quadrupedalism. Integrative and Comparative Biology. 2021;61:491–505.

28. Koprowski JL. Handling tree squirrels with a safe and efficient restraint. Wildlife Society Bulletin. 2002;30:101–3.

29. RStudio Team. RStudio: Integrated Development for R. RStudio, PBC, Boston, MA URL. 2021. http://www.rstudio.com/.

